# Deep genetic substructure within bonobos

**DOI:** 10.1101/2024.07.01.601523

**Authors:** Sojung Han, Cesare de Filippo, Genís Parra, Juan Ramon Meneu, Romain Laurent, Peter Frandsen, Christina Hvilsom, Ilan Gronau, Tomas Marques-Bonet, Martin Kuhlwilm, Aida M Andrés

## Abstract

Establishing the genetic and geographic structure of populations is fundamental both to understand their evolutionary past and preserve their future, especially for endangered species. Nevertheless, the patterns of genetic population structure are unknown for most endangered species, including some of our closest living relatives. This is the case of bonobos (*Pan paniscus*) which together with chimpanzees (*Pan troglodytes*) are humans’ closest living relatives. Chimpanzees live across equatorial Africa and are classified into four subspecies (Groves, 2001), with some genetic population substructure even within subspecies. Conversely, bonobos live exclusively in the Democratic Republic of Congo and are considered a homogeneous group with low genetic diversity (Fischer et al. 2011) despite some population structure inferred from mtDNA. Nevertheless, mtDNA aside, their genetic structure remains unknown, hampering our understanding of the species and conservation efforts. Placing bonobos’ genetics in space is however challenging because, being endangered, only non-invasive sampling is possible for wild individuals. Here, we jointly analyse the exomes and mtDNA from 20 wild-born bonobos, the whole-genomes of 10 captive bonobos and the mtDNA of 61 wild individuals. We identify three genetically distinct bonobo groups of inferred Central, Western and Far-Western geographic origin within the bonobo range. We estimate the split time between the central and western populations to ∼145,000 years ago, and genetic differentiation to be in the order of that of the closest chimpanzee subspecies. We identify putative signatures of differential genetic adaptation among populations for genes associated with homeostasis, metabolism and the nervous system. Furthermore, our estimated long-term *N*_*e*_ for Far-West (∼3,000) is among the lowest estimated for any great ape lineage. Our results highlight the need of attention to bonobo substructure, both in terms of research and conservation.

**Highlights:**

– We identified three genetically distinct populations of bonobos, inferred as having Central, Western and Far-Western geographic origin within the species range. The estimated split time is ∼145,000 years ago for the Central and Western populations, and ∼60,000 years ago for the two Western populations.
– The genetic differentiation between the Central and Far-Western bonobo populations is in the order of that between Central and Eastern chimpanzee subspecies, while the genetic differentiation among Western bonobo populations is similar to that among human groups.
– Once substructure is accounted for, we infer a long-term effective population size (*N*_*e*_) of only ∼3,000 for Far-Western bonobos, genetic isolation and inbreeding.

## RESULTS AND DISCUSSION

### Three genetically differentiated groups of bonobos

Bonobos are assessed as endangered on the IUCN Red List, with less than 20,000 extant wild individuals. Their recent historical range spans an area of 564,542 km^2^ (IUCN & ICCN, 2012), encompassing African forests within the Democratic Republic of Congo (DRC), where social unrest has limited research activities for decades (Whiten, 2020). Wild bonobos are thus less studied than other great apes. However, their biological similarity to humans, multi-male/multi-female societies, use of non-reproductive sexual behaviours, intergroup tolerance, and high social status of females make them uniquely interesting (Furuichi, 2011). Even though some cultural differences have been observed among communities (Hohmann & Fruth 2003, Samuni et al. 2020), from the point of view of their genomes, bonobos have long been considered a homogeneous group (Eriksson et al. 2004, Fischer et al. 2011) with low genetic diversity due to population bottlenecks (Eriksson et al. 2006, Prado-Martinez et al. 2013, de Manuel et al. 2016, Kuhlwilm et al. 2019). However, mitochondrial DNA (mtDNA) from seven wild communities revealed six haplogroups associated with specific geographic areas (Eriksson et al. 2006, Kawamoto et al. 2013), with the time to the most recent common ancestor inferred from the six mtDNA haplogroups being at least 380 thousand years ago (kya) (260-530 kya). MtDNA is a single locus and its genealogy reflects only the history of the maternal lineages, complicating the interpretation of inferences such as population split times. Since bonobos are primarily patrilocal, generating demographic inferences based on the nuclear genome is particularly important. Here, we use genomic datasets to assess the assumption of genetic homogeneity: the exomes (Teixeira et al. 2015) and mtDNA (Fischer et al. 2011) from 20 wild-born bonobos residing in an African sanctuary, together with 10 full genomes from wild born captive bonobos (Prado-Martinez et al. 2013) and the mtDNA of 136 wild individuals (Kawamoto et al. 2013).

A Principal Component Analysis (PCA) of the 20 exomes separates three distinct groups (Figure 1A), with two individuals (Lodja and Boende) falling in between groups. The clustering in separate groups is slightly higher than in 20 central chimpanzees (the chimpanzee subspecies with the highest genetic diversity; Prado-Martinez et al. 2013, de Manuel et al. 2016) and considerably higher than in 20 Yoruba humans (an African population with high genetic diversity, 1000 Genomes Project Consortium, 2010) (Figure S1A) that were sequenced using the same methodology (Methods). In fact, only bonobos show clear grouping of individuals in both PCs. We thus initially label these groups as B1, B2, and B3. The 20 exomes come from individuals of unknown geographic origin sampled in a sanctuary, so we can ensure that they are unrelated (Fischer et al. 2011, Teixeira et al. 2015) but are not able to assess sampling representation across the three groups. A neighbour-joining (NJ) phylogenetic tree separates B1-B2 from B3 (Figure 1B). An ADMIXTURE analysis (Figure 1C) shows *K*=3 as the best fit, and clustering of individuals in agreement with the PCA and NJ groups. Thus, PCA, NJ and ADMIXTURE indicate the presence of three distinct groups of bonobos in our 20 exomes.

**Figure 1.**
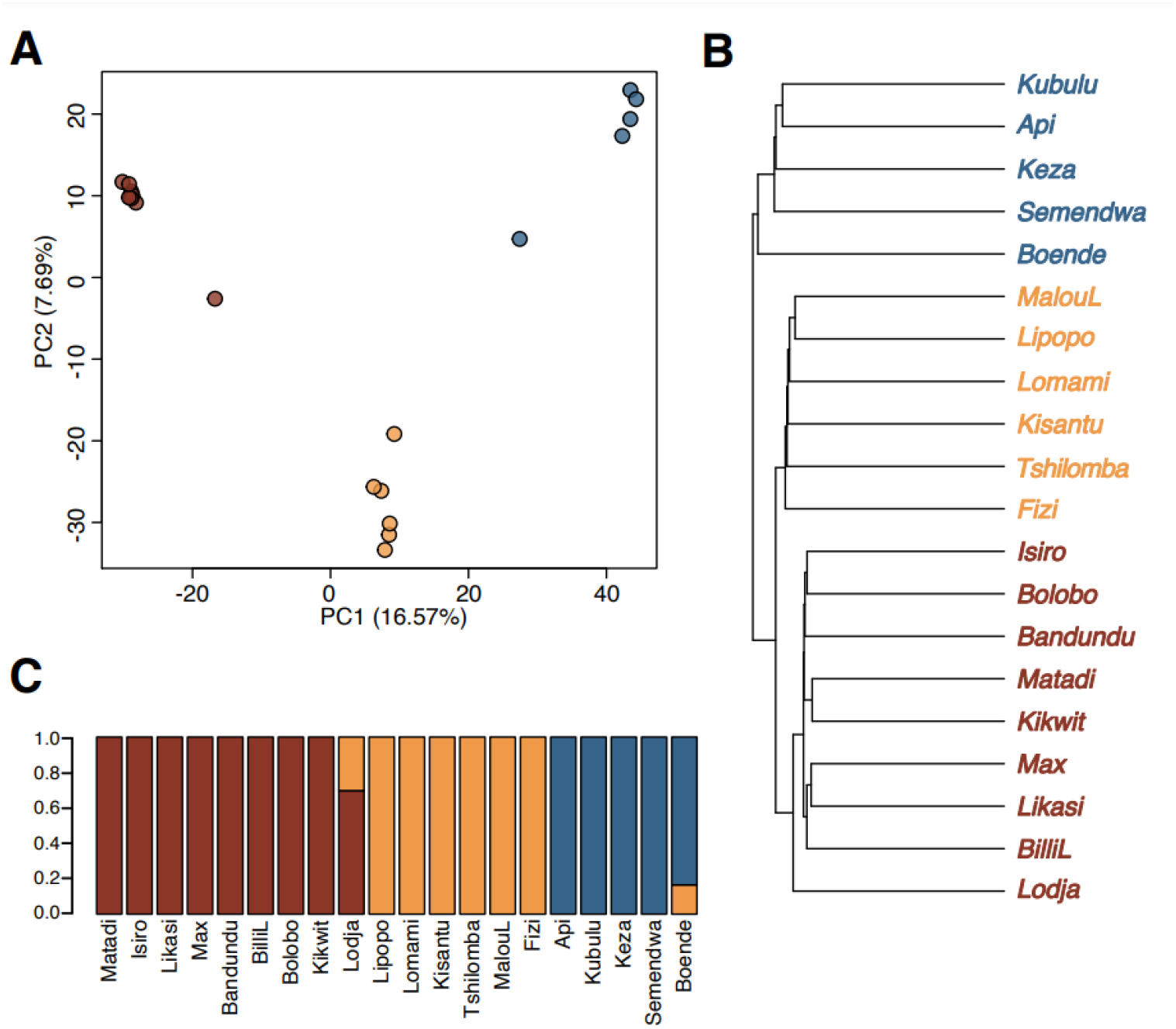
Three distinct groups of bonobos using whole exomes. **A**. PCA, **B**. NJ tree, with colours of groups according to PCA, and **C**. ADMIXTURE clustering of bonobos (k=3). Colours for the three groups are: B1 maroon, B2 orange and B3 blue. Comparable plots for chimpanzees and humans can be found in Figure S1A and Figure S1B. We note that the names of the individuals (as in B and C) do not necessarily reflect the place of origin.

We quantified the degree of population differentiation among the groups using pairwise F_ST_ (Table S1A). The highest differentiation is between B1-B3 (average F_ST_ = 0.145), followed by B1-B2 (F_ST_ = 0.093) and B2-B3 (F_ST_ = 0.088). To interpret these values, we put them in the context of humans and chimpanzees, the other great ape species for which we have fully comparable exome data. The F_ST_ values are in the range of those between chimpanzee subspecies: F_ST_ between B1-B3 is slightly higher than the F_ST_ between the closest chimpanzee subspecies, Central and Eastern, and slightly lower than between the other chimpanzee subspecies. Compared to human populations, the bonobo F_ST_ values are within the range between African and non-African humans (Table S1C). Of note, F_ST_ values do not directly reflect split times because other demographic factors that shape *N*_*e*_ (and thus drift) impact F_ST_. However, we suggest that the F_ST_ values reveal substantial genetic differentiation among the bonobo groups.

### The geographic origin of the three bonobo populations

Unfortunately, we lack precise information on the geographic origin of individuals from sanctuaries, hindering the geographic assignment of the three inferred groups (B1-B3). Nevertheless, we were able to take advantage of 61 mtDNA haplotypes from seven wild bonobo populations of known location (Kawamoto et al. 2013 and Figure 2A) to make inferences about the geographic origin of our samples. The populations are not completely differentiated with a few haplotypes from the Central populations and the Eastern population close to the Western group, in a multidimensional scaling (MDS), likely due to female migration and/or higher effective population size of Central populations. However, the mtDNA sequences of the 20 sanctuary bonobos fall perfectly within the MDS clusters of the Kawamoto mtDNA sequences (Figure 2A). Since the Kawamoto samples are placed geographically, and there is a strong correlation between autosomal and mitochondrial pairwise nucleotide distances (Mantel test based on Spearman rank correlations = 0.73; *p-value* = 0.001), this analysis provides the likely geographic origin of the bonobos in the B1, B2 and B3 groups.

**Figure 2.**
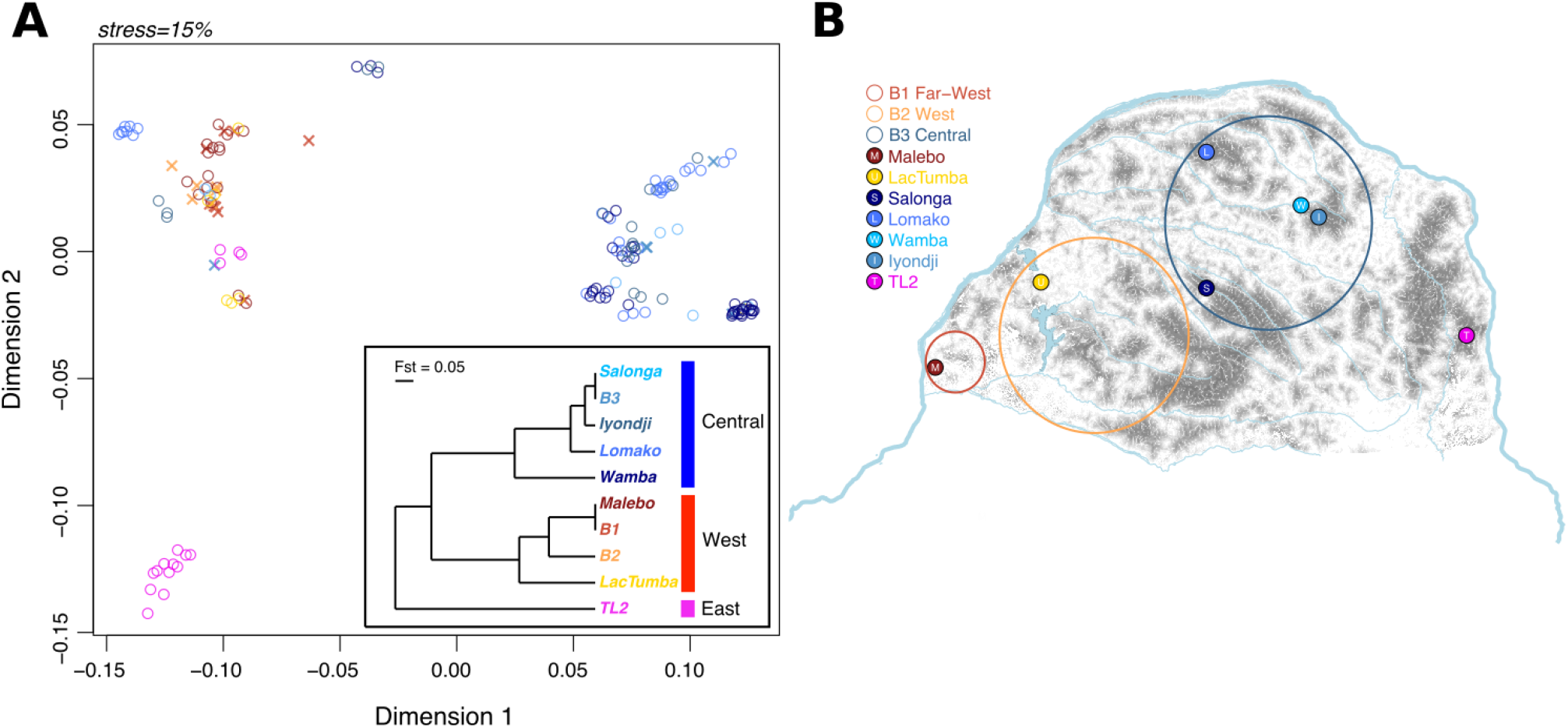
**A**. MDS of bonobo individuals, combining the mtDNA sequences from our samples (x symbols), published in Fischer et al. 2011, and from Kawamoto et al. 2013 (circles). For a better visualisation we added jittering (see Figure S2 without jittering and the Methods). Inset: UPGMA tree computed for the ten different sample groups using Φ_ST_ (Michalakis & Excoffier, 1996). **B**. Sampling location of the bonobo populations from Kawamoto et al. (2013) in filled circles and a potential area of origin of our inferred populations in empty circles. The gray areas reflect the probability of suitable habitat for bonobos according to Hickey et al. (2013), where the darker the area the higher the probability. The main rivers and lakes of the Democratic Republic of Congo are shown in light blue.

In a phylogenetic tree, the Kawamoto mtDNA samples show geographic substructure, with a western, a central and an eastern clade (Kawamoto et al. 2013). Incorporating the B1, B2 and B3 groups in this mtDNA tree does not affect its topology. In agreement with the PCA, NJ and F_ST_ results, both B1 and B2 fall within the same cluster, the western clade (Figure 2A). B3 falls in a separate cluster, the central clade, and has mtDNA Φ_ST_=0 when compared to Salonga and Iyondji, suggesting that B3 is closely related to these populations. Interestingly, B1 clusters with the Malebo population (Φ_ST_=0), which lives at the western periphery of the bonobo range and is characterised by low levels of mtDNA nucleotide diversity and higher genetic distance to the other western community LacTumba. Hereafter, we thus refer to the B1 group as the ‘Far-West’ population, B2 as the ‘West’ population, and B3 as the ‘Central’ population, based on their likely geographic distribution according to these genetic relationships, and to the Far-West and West populations together as the Western group. None of our samples falls within the eastern group (namely TL2 in Kawamoto et al. 2013), so we are unable to make inferences about this population. Based on this analysis together with geographical nature within the bonobo habitat, a likely scenario of the geographic range of the three bonobo populations is shown in Figure 2B.

### An old split between the western and the central populations

The high level of genetic differentiation among bonobo populations makes estimating split times interesting. Such estimates are most accurate when based on whole-genomes, where many neutral loci can be analysed, so we used ten published high-coverage whole-genomes (Prado-Martinez et al. 2013). First, to explore the genetic relationships between these and the sanctuary samples, we extracted the exomes of the Prado-Martinez genomes and combined them with the 20 exomes. In a PCA analysis of these 30 exomes, the ten Prado-Martinez individuals cluster well with the three populations we identified (Figure S3). Hence, we used one genome chosen randomly from each population to make genome-wide inferences about the Far-West, West and Central populations. We used G-PhoCS (Gronau et al. 2011), which infers the demographic parameters such as population divergence times, effective population sizes, and migration rates, by analysing unlinked, short, putatively neutral loci. G-PhoCS is a suitable method when sample size per population is small and restricted to a single individual per lineage, as in this study.

For the G-PhoCS analysis, we assumed a phylogenetic tree with the three bonobo populations identified and a western chimpanzee as an outgroup, allowing migration among bonobo populations shortly after their split time (Figure S3B). We chose a western chimpanzee as an outgroup as previous evidence suggests that this subspecies has not received direct gene flow from bonobos (de Manuel et al. 2016). We infer low levels of gene flow among the groups (Figures 3, S4-S6, Table 1), while confirming that they belong to discrete populations. The estimated split time between Central and Western bonobos is ∼145 kya (95% Bayesian credible interval (CI), 116-302 kya; see Table 1), and between West and Far-West ∼60 kya (95% CI, 23-116 kya), whereas between chimpanzees and bonobos it is ∼1.29 million years ago (mya) (95% CI, 1.21-1.37 mya). Performing the analysis multiple times, CIs overlap between runs, reflecting high concordance among G-PhoCS runs (see Methods and Figure S4). As always, such inferences should be considered approximate estimates of population split times, because population splits are rarely instantaneous, being often rather complex. For context, the estimated split time between the Central and Western bonobo populations is close to the estimated split time between the Central and Eastern chimpanzee subspecies (∼139 kya, de Manuel et al. 2016). The estimated split time between the two Western populations, on the other hand, is closer to the inferred split time between East African and non-African human populations (65,000 years ago; Li & Durbin, 2011, Schiffels & Durbin, 2014, Hershkovitz et al. 2015).

**Table 1.** Estimate summaries for each parameter across 20 G-PhoCs runs using 900 MCMC samples per run. Mean and 95% Bayesian credible intervals (Cis) of split times (T), effective population sizes (Ne) and migration probabilities (m) between populations.

| Parameters | with Migration |  |  | without Migration |  |  | Units |
| --- | --- | --- | --- | --- | --- | --- | --- |
|  | estimate | low | high | estimate | low | high |  |
| T_farWest-West | 59.93 | 23.26 | 116.28 | 36.76 | 23.26 | 46.51 | kya |
| T_Western-Central | 144.66 | 116.28 | 302.33 | 127.46 | 116.28 | 139.53 | kya |
| T_Bonobo-Chimp | 1289.20 | 1209.30 | 1372.09 | 1198.44 | 1139.53 | 1279.07 | kya |
| Ne_farWest | 2.84 | 1.60 | 4.01 | 2.64 | 1.60 | 4.01 | k individuals |
| Ne_West | 4.55 | 1.60 | 7.22 | 3.69 | 2.41 | 5.61 | k individuals |
| Ne_Central | 6.84 | 4.81 | 8.02 | 7.53 | 7.22 | 8.02 | k individuals |
| Ne_Chimp | 12.59 | 12.03 | 13.63 | 13.43 | 12.83 | 14.43 | k individuals |
| Ne_Western | 28.68 | 8.82 | 65.76 | 26.33 | 17.64 | 37.69 | k individuals |
| Ne_Bonobo | 23.23 | 21.63 | 24.86 | 22.28 | 20.85 | 23.26 | k individuals |
| Ne_Ancestral | 217.00 | 211.71 | 222.13 | 218.34 | 213.31 | 223.74 | k individuals |
| m_farWest-West | 0.199 | 0.007 | 0.497 | NA | NA | NA | probability |
| m_Western-Central | 0.222 | 0.000 | 0.979 | NA | NA | NA | probability |
| m_Bonobo-Chimp | 0.051 | 0.027 | 0.085 | NA | NA | NA | probability |

**Figure 3.**
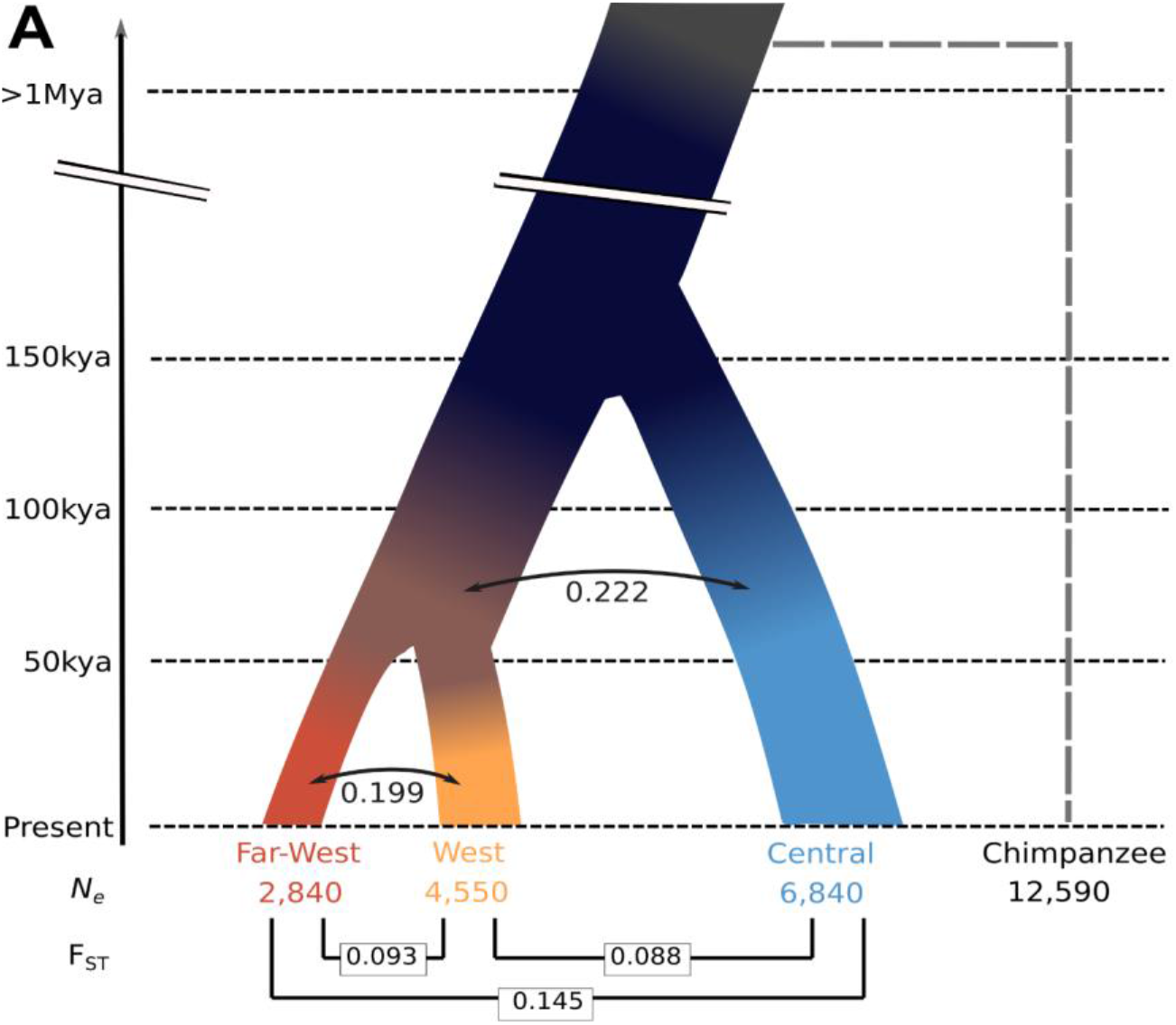
Conceptual model of split times and effective population sizes (*N*_*e*_) for the three bonobo populations, as well as migration probabilities, as estimated by G-PhoCS. One western chimpanzee individual was used as an outgroup. kya, thousand years ago; Mya, million years ago. The values here point estimates, with details and Bayesian CIs in Table 1. F_ST_ values are noted at the bottom. Colors for the three groups are: maroon for Far-West, orange for West and blue for Central

We calculated ‘mutual migration probability’ as a measure of gene flow based on the parameters inferred by G-PhoCS (Methods) that indicates the probability that two populations exchanged genetic material, irrespectively of the direction, since they diverged and until the next population split. The calculated probabilities of migration across the 20 replicate runs appear to fluctuate more across runs than other demographic parameters such as split time and *N*_*e*_. Migration probabilities estimated between West and Far-West populations in particular show high variation across replicates (Figure S4). This is likely due to the algorithm being unable to differentiate between deeper divergence with frequent introgression, or more recent divergence with sporadic introgression, as shown by the positive correlation between estimated migration probabilities and estimated divergence times (Figure S6). As expected, a model without migration (Figure S3B) provides similar albeit slightly younger estimates: the split time between the Central and Western populations is estimated to ∼127 kya (95% CI, 116-140 kya; see Table 1), between the West and Far-West populations to ∼37 kya (95% CI, 23-47 kya), and between chimpanzees and bonobos to ∼1.20 mya (95% CI, 1.14-1.28 mya).

### Different genetic diversity, effective population size (*N*_*e*_) and inbreeding

The levels of genetic diversity differ substantially among populations (Fig S7D), with the mean number of pairwise differences in Central being almost 1.4 times that in Far-West (Far-West has 3.28, West 4.03, and Central 4.53 in windows of 10,000 base pairs [kb]). Heterozygosity (Figure S7C) shows similar patterns (Far-West has on average 2.16, West 2.59, and Central 2.56 heterozygous sites in 10 kb), and both measures agree well with mtDNA haplotype diversity (Kawamoto et al. 2013), where central populations show higher levels of mtDNA diversity than western populations. Differences in genetic diversity often reflect differences in demographic history. G-PhoCS’s estimated *N*_*e*_ is lowest for the Far-West bonobos (2,840, 95% CI, 1,600-4,010) compared to West (4,550, 95% CI, 1,600-7,220) and Central (6,840, 95% CI, 4,810-8,020) (Figures 3, S4, S5 and Table 1). The estimated Central *N*_*e*_ is more than two times larger than the Far-Western *N*_*e*_, suggesting substantial differences in the demographic history of the populations. Further, some of these *N*_*e*_ estimates are even lower than previously reported for bonobos as a whole, which range from 5,000 to 29,129 (Prado-Martinez *et al*. 2013; Kuhlwilm *et al*. 2019). This is not surprising, as these were estimates of a long-term *N*_*e*_ for bonobos as a species, while we are estimating its specific populations since their divergences. However, it is notable that the *N*_*e*_ estimate of Far-West bonobos is among the lowest *N*_*e*_ estimates across all great ape groups, comparable only to Mountain gorillas and Tapanuli Orangutans with ∼2,200 (Pawar et al. 2023) and ∼2,500 (Nater et al. 2017), respectively. Of note, the estimated *N*_*e*_ of the common ancestor of the three bonobo populations is 23,230 (Table 1), which is in line with the estimated *N*_*e*_ of bonobos from Kuhlwilm et al. (2019).

A possible effect of small populations is inbreeding, the mating of closely related individuals. Inbreeding increases homozygosity, which can expose deleterious recessive alleles in inbred populations. To investigate putative recent inbreeding, we examined the distribution of runs of homozygosity (ROH), which are continuous segments depleted of heterozygous positions in an individual due to a shared recent ancestor. Bonobos as a group have on average longer ROHs than our samples of humans and chimpanzees, as expected from their low *N*_*e*_ (Figure S7A). Among bonobos, the Far-West individuals have the longest ROHs, which are on average 29% longer than in the West and the Central bonobos. Strikingly, these are 61% longer than in humans and 139% than in chimpanzees (Figure S7A). Not surprisingly, Far-Western bonobos also have the highest level of identity-by-descent (IBD) segments. IBD segments are inferred identical chromosome fragments between two individuals sharing the same recent common ancestor, resulting from mating in small and/or inbred populations. All bonobos have IBD segments, shared predominantly with other members of their population, as is the case in our samples of chimpanzees and humans (Yoruba). Still, bonobos in the Far-West show longer IBD segments than bonobos in the other populations, with a total IBD length twice as long as the other two populations (Figure S7A). Far-West bonobos are also related to a much larger extent than Central and even West populations, which suggests that they are more inbred (Figure S7B). G-PhoCS infers migration between Central and the common ancestor of the Western bonobos, and migration between West and Far-West populations (mutual migration probability of 0.22 and 0.2, respectively, Table 1). Yet our results suggest that even in the presence of some gene flow, the Far-West is genetically differentiated from the other populations, likely as a result of their fragmented habitat (see the area surrounding the Malebo population in Figure 2B).

### Signatures of positive selection

Genetic adaptation has contributed to the genetic differentiation among chimpanzee subspecies, as shown by analyses of whole genomes from these four subspecies (Cagan et al. 2016; Schmidt et al. 2020; Pawar et al. 2022). The bonobo populations we identified here may also have adapted to their local environment, since they are genetically distinct and geographically differentiated. We explored potential differential adaptations among bonobo populations with GRoSS (Refoyo-Martinez et al. 2019), which detects unusually large allele frequency changes generated by positive selection. We analysed every SNP in the exome and averaged their *p*-values within each exon, to obtain an exon-based distribution of GRoSS empirical *p*-values as potential signatures of local adaptation. While this empirical approach does not demonstrate that local adaptation among bonobo populations has taken place, it allows us to identify the genes with the strongest allele frequency change which are the most likely targets of putative local adaptation. As expected, exons with the lowest *p*-value differentiate the three populations in a PCA (Figure S8A) and their average difference in allele frequency between populations is higher than in the rest of the exome (0.21-0.27 in outlier exons vs. 0.121-0.16 exome-wide for Far-West, 0.174-0.193 vs 0.121-0.146 exome-wide for West, 0.296-0.251 vs 0.16-0.146 exome-wide for Central). However, these SNPs are not strongly differentiated, suggesting that if there is local adaptation, it is not monogenic. Only 17 exons have a GRoSS *p*-value < 0.01 after multiple testing correction (Benjamini-Hochberg): 11 in Far-West, four in West and two in Central. Larger sample sizes are needed to infer selection signatures with confidence, but these are potential candidates to mediate population-specific adaptation.

The most significant exon for Far-West is in *SCNN1G* (adjusted GRoSS *p*-value = 0.0016, Figure S8B), a gene that encodes for a sodium channel subunit involved in electrolyte homeostasis (Strautnieks et al. 1996). Far-West-specific changes in this gene do not contain protein-coding changes, but one exon contains four potentially regulatory variants that differ in frequency 60% to 100% between Far-West and the other populations (Table S2). The transcription factor-coding gene *PRDM10* contains one significant exon (adjusted GRoSS *p*-value = 0.0020) that includes a missense variant (P>T) with 86% higher frequency in Far-West compared to the other populations. This gene shows signatures of positive selection in the ancestor of modern humans after the split from Neandertals (Peyregne et al. 2017, Kuhlwilm & Boeckx, 2019). In West, the most significant exon is in *CYB5R2* (adjusted *p*-value = 0.0023), which encodes for an enzyme involved in cholesterol biosynthesis and carries several non-coding SNPs fixed in this population and absent in the other two populations (Table S2). In Central, the highest-ranking exon is *GABRE* (adjusted *p*-value = 0.0035), a gamma-aminobutyric acid A receptor with various functions in the central nervous system. Although these results should all be taken with caution, the presence of significant exons suggests that the different populations may have been subject to differential selection pressures that generated genetic differentiation among populations of bonobos.

## Conclusions

Our results reveal the presence of strong population substructure in bonobos, indicating at least three distinct populations in the west and the central part of their range. The genetic differentiation and estimated divergence time between the Central and Western populations is of comparable depth to those between the closest chimpanzee subspecies, Central and Eastern. Bonobos live exclusively in the DRC and although they have been studied for some time, thus far little behavioural diversity has been observed among groups, with only some differences reported in drumming, grooming and hunting among central bonobo communities– the best-studied ones through long-term data collection (Hohmann & Fruth 2003, Samuni et al. 2020). It is notable that the three bonobo populations are not only genetically differentiated, but they also have substantially different evolutionary histories, have experienced different levels of inbreeding, and might be differentially adapted to their local environment. The Far-West populations show evidence of particularly long-term low *N*_*e*_ and genetic isolation (with sporadic gene-flow), making them potentially vulnerable to future ecological changes. Of note, our sampling does not appear to cover the eastern population (TL2 in Figure 2A), which previous mtDNA studies identified as potentially highly differentiated (Kawamoto et al. 2013, Takemoto et al. 2017). Thus, our work stresses the importance of performing genomic studies of wild bonobos across their full geographical range. More generally, the presence of genetically differentiated bonobo populations, and the high genetic isolation of some of these populations, should be taken into consideration in the planning of conservation efforts.

## Materials and Methods

### Data Preparation

We analysed whole-exome high-coverage (∼20X) Illumina sequencing data of 20 bonobos (*Pan paniscus*), 20 central chimpanzees (*Pan troglodytes troglodytes*) and 20 humans (*Homo sapiens*) that we published previously in Teixeira et al. 2015. Bonobo and chimpanzee blood samples were collected in African sanctuaries (Lola ya bonobo sanctuary in Kinshasa, DRC; and Tchimpounga sanctuary, Jane Goodall Institute, Republic of Congo, respectively) and human samples belong to the Yoruba population from HapMap (Teixeira et al. 2015). The mtDNA sequences of the 20 bonobos were previously published (Fischer et al. 2011). We also used 10 high-coverage (∼27X) bonobo whole-genomes which are from wild-born individuals residing in European zoos (Prado-Martinez et al. 2013). The protocols on how exomes and whole-genomes were mapped and prepared are available in these previous publications (Teixeira et al. 2015, Prado-Martinez et al. 2013).

### Population substructure and differentiation

PCA were performed using the function ‘glPca’ from the R-package “adegenet” (Jombart 2008) and run for all individuals together and separately per species. NJ trees were generated using the function ‘nj’ from the R-package “ape” (http://ape-package.ird.fr/) also constructed for all individuals together, and separately per species, with a distance matrix of pairwise nucleotide differences such that the distance between a heterozygote and any homozygote is 1, and the distance between the two different homozygotes is 2. This was used to calculate the number of pairwise differences between individuals, which was then divided by the total number of base-pairs that passed the filters in all species (25,781,213) to have a comparative measurement. The number of variant sites across all autosomal sites (SNPs and/or fixed differences) is 228,488 for chimpanzees, 86,250 for bonobos, and 106,832 for Yoruba humans. ADMIXTURE (Alexander, Novembre, & Lange, 2009) was run on a subset of SNPs that minimizes linkage disequilibrium (LD) by removing high-LD SNPs with plink (Purcell et al. 2007) with the following steps: 1) create a window of 200 SNPs; 2) calculate LD (as r^2^) between each pair of SNPs in the window; 3) if r^2^ > 0.5 remove one of a pair of SNPs; 4) shift the window 20 SNPs and repeat the procedure. ADMIXTURE was run from K=1 to K=8, each with 10 replicates using the following command line: “admixture -s time --cv INPUT.ped k”. We run the cross-validation procedure (“--cv” flag) as described in (Alexander et al. 2009) to determine the number of K that best fits the data. The software CLUMPP (Jakobsson & Rosenberg, 2007) was used to condense the 10 admixture runs per K to identify modes where different runs have similar outcomes (>90%) by selecting pairs of replicates having a symmetric similarity coefficient G‘ > 0.9. To measure pairwise population differentiation, we calculated the average F_ST_ across all sites that are polymorphic in at least one of the two populations, using Weir and Cockerham’s F_ST_ (Weir & Cockerham, 1984). The pairwise comparisons are: 1) among bonobo groups using our exome data; 2) among all chimpanzee subspecies using the GAGDP dataset, and among two central chimpanzee subgroups identified with our exome data; 3) among six human populations from Africa (Yoruba and Luhya), Europe (Toscani and Finns) and Asia (Han Chinese and Japanese) using the 1000 Genomes data (1000 Genomes Project Consortium, 2010). Heterozygosity is calculated for each individual simply as the number of heterozygous sites over the total number of base pairs.

### mtDNA analyses

Kawamoto et al. 2013 analysed the D-loop of the mtDNA of 136 bonobo individuals to generate a phylogenetic tree of samples of known geographic origin. Together with the published mtDNA data (Fischer et al. 2011) of our 20 bonobos, in total 156 mtDNA sequences were aligned with the software ‘mafft’ v7 (Katoh & Standley, 2013), and filtered to remove positions with indels and missing data, retaining 1,101bp. We calculated pairwise sequence differences using the Kimura 2-parameters model (Kimura, 1980) and performed MDS on these distances using the R-function ‘cmdscale’. MDS is a dimensional reduction, similar to PCA albeit using different methodologies, which allows us to visualize the information contained in a matrix of pairwise distances with some degree of loss function called “stress”. In order to visualize samples with identical sequences/haplotypes, we randomly added noise (or jittered) as 10% of the standard deviation for each dimension of the MDS, which are 0.010 and 0.0045, for Dimensions 1 and 2, respectively. We calculated Φ_ST_ among 10 bonobo groups according to the formula of (Michalakis & Excoffier, 1996), which then was used for building an UPGMA tree. The result without jittering is plotted in Figure S2. For plotting, we used the function ‘upgma’ of the R-package ‘phangorn’ (Schliep, 2011) with default parameters. To build NJ-trees, as for the autosome, we used the function ‘nj’ from R-package ‘ape’ (http://ape-package.ird.fr/).

### Demographic inference

To infer the demographic history of the three populations, we used the Generalized Phylogenetic Coalescent Sampler (G-PhoCS; Gronau et al. 2011), a Bayesian sampling method that summarizes the information over local genealogies at short, putatively neutral loci in approximate linkage equilibrium. G-PhoCS infers demographic parameters, such as divergence times, effective population sizes and migration rates, given a pre-specified population phylogeny. We used the UPGMA tree from their mtDNA haplotypes (Figure 2A), which is consistent with the NJ tree topology from autosomal data (Figure 1B), for the tree topology of three bonobo populations and a western chimpanzee (Figure S3B). G-PhoCS can produce reliable parameters using one full genome per population, so we used bonobo whole genomes from Prado-Martinez et al. 2013, one genome per bonobo population, which were selected based on their PCA clustering with the exomes (Results and Figure S3A). Our analysis follows best-practice procedures previously defined (Gronau et al. 2011; Kuhlwilm et al. 2016; Schlebusch et al. 2017), including filters and parameters. We used eight quality filters downloaded from the UCSC genome annotation database for hg19 (http://hgdownload.cse.ucsc.edu/goldenpath/hg19/database/, the last date of access 29/04/2019) to remove known genic regions (refGene, knownGene), simple and complex repeat regions (simpleRepeat, genomicSuperDups), CpG islands (cpgIslandExt), repeat masker (rmsk), conserved regions across 46 placental species (phastConsElements46wayPlacental), and synteny net between the assemblies hg19 and PanTro4 (netPanTro4). We computed a set of 1000 bp long loci that avoided the genomic filters and separated by at least 10 kb. This resulted in 72,607 loci overall. We then partitioned this set into three subsets of 24,202 or 24,203 alternating loci, and used the first two subsets in our analysis (subset 1 and 2). Finally, we removed loci with more than 20% missing data for one of the analysed individuals. As a result, each analysis considered 23,439 and 23,424 independent loci sets that are separated by at least 30 kb. These properties were shown to minimize the impact of recombination within loci and maximize independence between loci (Gronau et al. 2011). Only fragments with less than 20% missing data within each individual were analysed. The numbers of fragments used in the G-PhoCS run are 23,439 for Set 1 and 23,424 for Set 2. We ran G-PhoCS with and without migration (Figure S3B), in both cases with 1,000,000 MCMC iterations in each run. We ran 10 replicate runs using each loci set and in each migration scenario, which results in 20 replicates per migration scenario. To estimate split time (Tau) and *N*_*e*_ (Theta), we discarded the first 100,000 iterations as burn-ins, taking one estimate every 1,000 iterations in order to minimize the issue of auto-correlation, which yielded 18,000 MCMC samples per parameter in each migration scenario. We then calculated mean values as point estimates and 95% Bayesian credible intervals as plausible ranges. To convert Tau and Theta to generations and *N*_*e*_, we used a mutation rate of 0.43 × 10-9 per site per generation (Besenbacher et al. 2019). We further used 29 years as an average generation time to have the split times in calendar years. For migration between populations, we use ‘migration probability’, which ranges between 0 and 1, and calculated as P = 1 — *e*^-rate^. This is a total rate, which is a product of the migration rate and time of migration, summing the rates in both directions.

### IBS, IBD and ROH

Identity-by-state (IBS) can be observed at a given locus, for any given pair of individuals with genotype information, with three possible outcomes: the individuals have two different alleles (IBS0) or they share one (IBS1) or two (IBS2) alleles. Two individuals who share 1 or 2 alleles IBS at a given locus may have inherited them from a recent common ancestor, in which case these alleles are identical-by-descent (IBD). We inferred IBD using a software KING (Manichaikul et al. 2010) to infer the length and the number of IBD segments. Importantly, KING design allows robust inference of IBD segments in the presence of population substructure, as is the case in the populations used in this study. We ran KING with the “--ibdseg” and “--related” options, and other parameters were set to default values. IBD regions tend to be short between pairs of individuals derived from a given population that are not closely related, primarily because their last common ancestor was many generations ago; they tend to be long among closely related individuals. Using the set of informative nucleotide positions for the three species, we computed the number and length of the IBD regions (defined here as regions between IBS0 alleles) among all the individuals. ROH, which are continuous segments depleted of heterozygous positions likely because the two chromosomes, are derived from the same recent ancestor. As exome data are composed mostly of the genic part of the genome and cannot be used to infer with precision the ends of ROHs, we restricted the analysis to the comparison among individuals. We then compared the average ROH value in each bonobo group with the value of central chimpanzees and the value of Yoruba humans sequenced in the identical way.

### Positive selection

To identify potential targets of differential adaptation among populations, we applied GroSS (Refoyo-Martinez et al. 2019). This method uses allele frequency data to identify genomic regions putatively under positive selection, and the branch where selection took place. Here, the three bonobo populations can be represented by a simple three-branch tree (Fig. 3), where Far-West and West split from the common ancestor with Central. We note that the method does not distinguish between selection on the common branch of Far-West and West, or on the Central branch. For each individual, we removed sites with less than 5 reads, more than 99 reads, or genotype quality below 20, resulting in 1,078,889 segregating sites, and ran GroSS (https://github.com/FerRacimo/GroSS) using R version 3.4.2. We processed the output in the R environment, averaging the raw p-values for each exon with more than 3 SNPs (39,133 exons). Few exons reach nominal significance across the genome (at a p-value of < 0.01: Far-West: 36; West: 24; Central: 23). We assessed the most significant exons in a PCA (using an adjusted p-value cutoff of < 0.05) for each of the three branches (Far-West: 93 exons; West: 74 exons; Central: 75 exons), using the R-package ‘adegenet’ (Jombard & Ahmet, 2011). We inferred functional consequences of the genetic variants in these exons using the Ensembl Variant Effect Predictor (McLaren et al. 2016).

## Acknowledgments

C.d.F and A.A were supported by intramural funds from the Max Planck Institute for Evolutionary Anthropology (Leipzig, Germany). CdF was also supported by grant agreement no. 694707 “100 Archaic Genomes” assigned to Svante Pääbo. AA was additionally funded by the Wellcome Trust ISSF3 UCL award 204841/Z/16/Z. M.K. was supported by “la Caixa” Foundation (ID 100010434), fellowship code LCF/BQ/PR19/11700002, the Vienna Science and Technology Fund (WWTF) [10.47379/VRG20001] and by the Austrian Science Fund (FWF) [FW547002]. We would like to thank Svante Pääbo for access to samples and discussions, Mimi Arandjelovic and Martin Surbeck for their valuable comments, Roger Mundry for the R-programming discussions, Hjalmar Kühl for sharing the data of the bonobo’s habitat and Hiroyuki Takemoto for providing the location of the sampled individuals analysed for mtDNA. We are grateful to the Life Science Compute Cluster of the University of Vienna.

## Author contributions

A.A. and CdF conceived the project. S.H., C.d.F., G.P., J.R.M., R.L., M.K. and P.F. analysed data, with support from A.A., I.G., S.P., T.M.B and C.V. A.A., T.M.B. and M.K. provided funding. S.H., C.d.F and A.A. wrote the manuscript with input from all authors.

