## Supplementary for "Deep genetic substructure within bonobos"

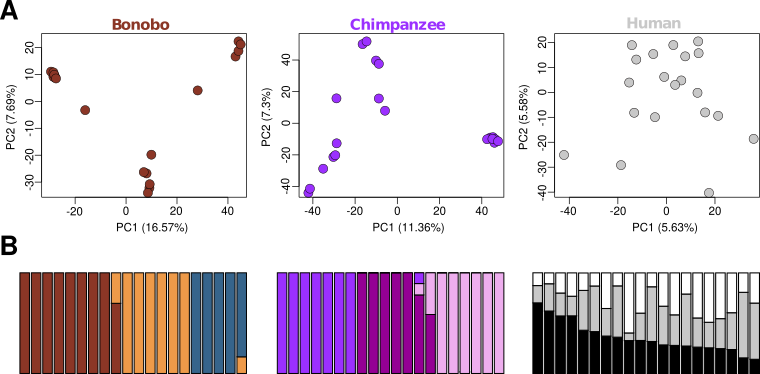

**Figure S1.**

**A.** PCA results of the analysis of each species separately, using 20 exomes per species: bonobos, central chimpanzees and Yoruba humans. We observe three groups in bonobos, two groups in chimpanzees, and no grouping in humans. **B.** ADMIXTURE results for bonobo, chimpanzee and human, K=3.

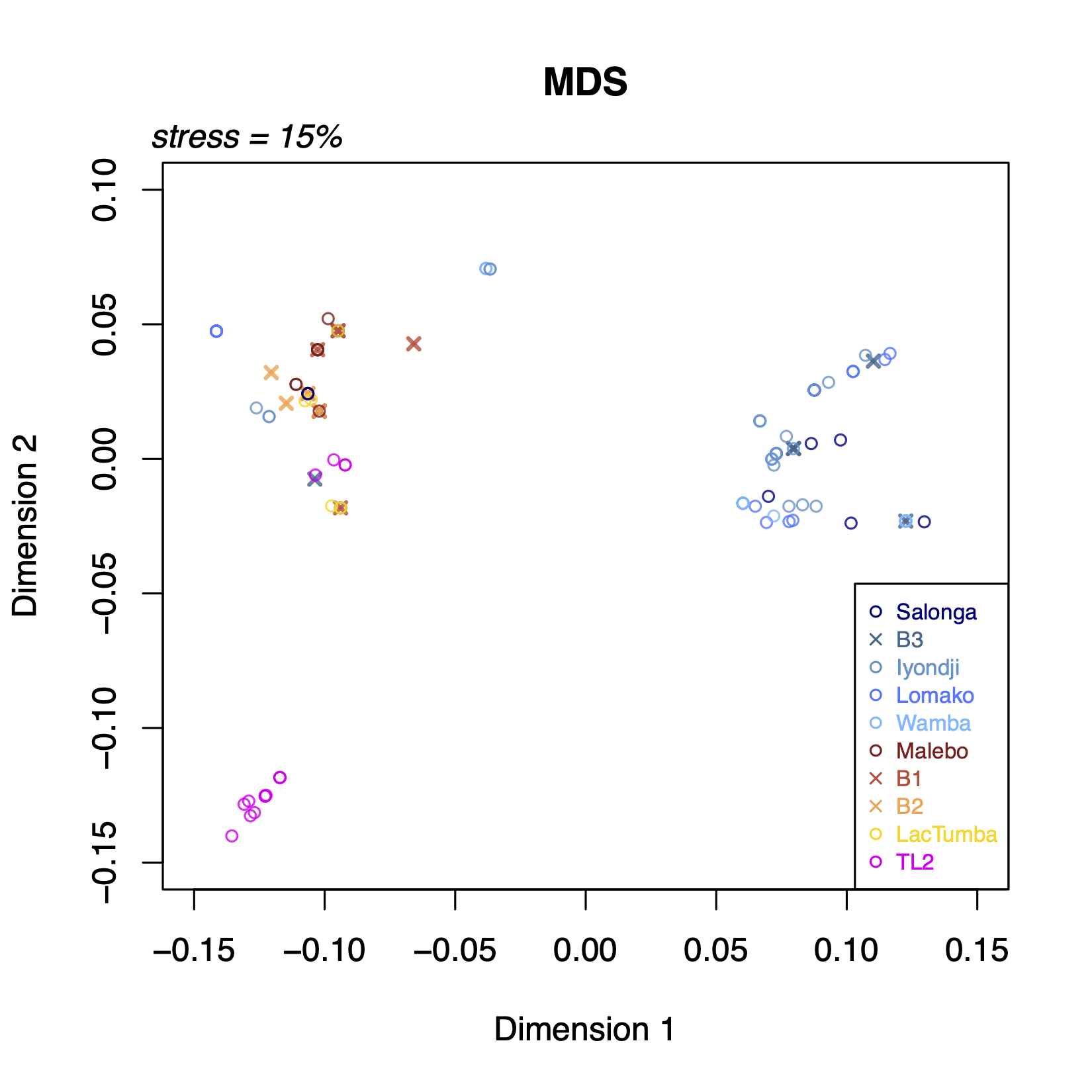

**Figure S2**

Multidimensional scaling (MDS) of bonobo individuals, combining the mtDNA sequences from our samples (x symbols), published in Fischer et al. 2011, and from Kawamoto et al. 2013 (circles). This is the same as Figure 2 but without jittering the same haplotypes.

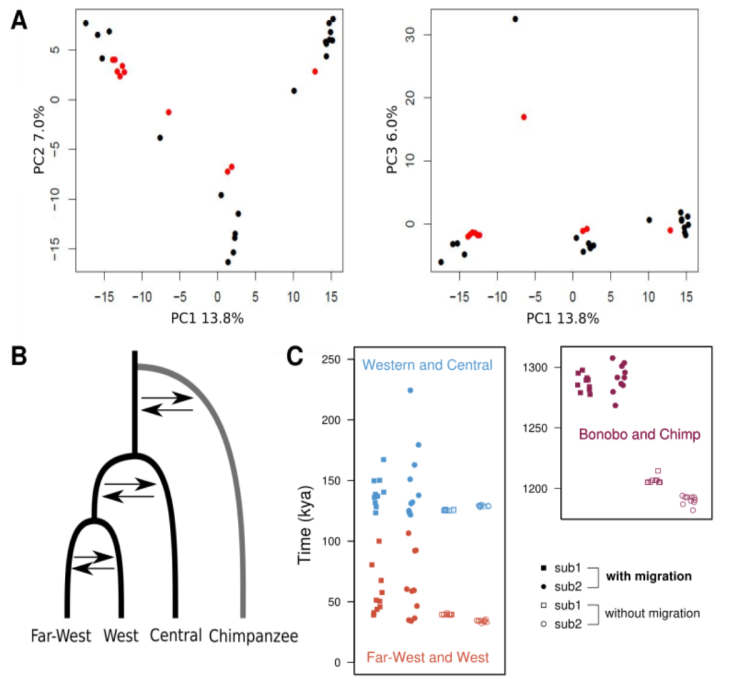

**Figure S3**

**A.** PCA results of the 30 exomes: 20 exomes (Teixeira *et al.* 2015) in black and 10 exomes (Prado-Martinez *et al.* 2013) in red. **B.** Schematic representation of the phylogenetic tree used for G-PhoCS, where all three bonobo populations and a Western chimpanzee lineage are nested and with migration bands for the G-PhoCS model with migration. **C.** The distribution of G-PhoCS estimates for split times from the twenty replicate runs (ten for each subset, sub1 and sub2, see Methods and also figures S4-S6), in the models with and without migration. Each split node is coloured according to the legend.

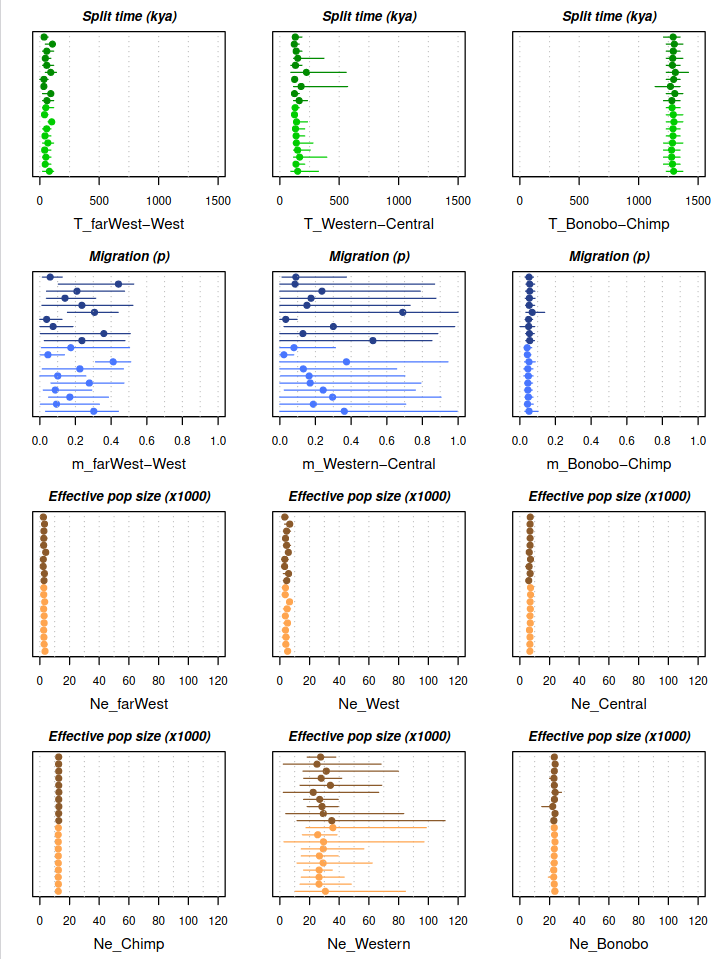

**Figure S4**

Estimates across 20 runs for each parameter. The dots represent the mean of 900 MCMC samples, i.e. one run (see Materials and Methods) and the horizontal lines the 95% quantile of these values. In green are the split times, in blue the migration probabilities and in orange the effective population sizes. Within each parameter, in light and dark colors are the 10 runs for subset 1 and subset 2, respectively.

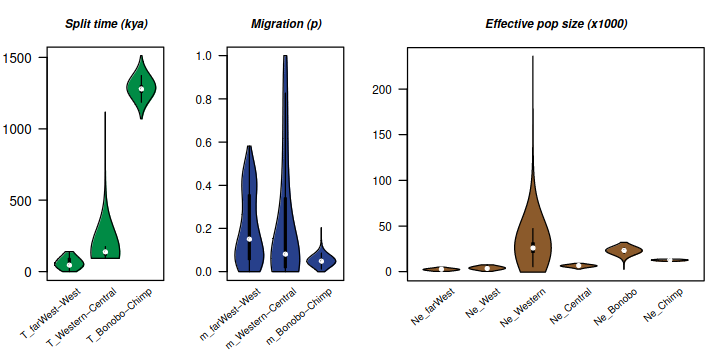

**Figure S5**

Distribution of the 18000 MCMC samples for each parameter across the 20 G-PhoCS runs with migration. The violin plot has been generated using the function ‘vioplot’ with default parameters (except for the style) from the R-package ‘vioplot’. The white dots represent the median of the distributions, while for the estimates reported in Table 1 we used the mean.

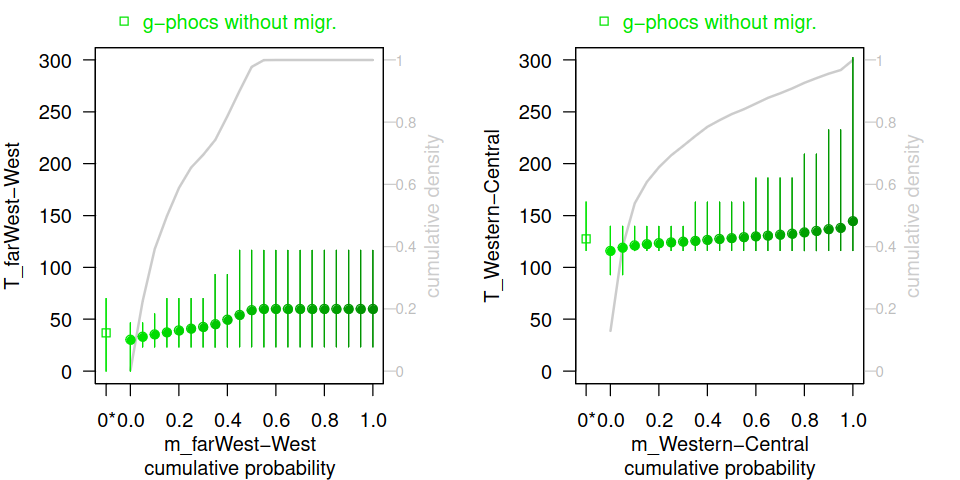

**Figure S6**

Split times stratified by migration. The figures show the split time estimates (y-axis) using different threshold values for the migration probability (x-axis). We used 21 bins for the cumulative density distribution of the migration probability (depicted with gray lines and in the y-axis on the right its cumulative density). The full circles represent the estimates and the vertical bars the entire range of the values, while the empty squares represent the estimates and the entire range when using the G-PhoCs runs without allowing migration (indicated also with ‘0*’ on the x-axis). For an easier view, we colored the dots according to the migration probabilities: darker the color higher the probabilities. The plot on the left is for the split time between the two Western populations (T_farWest-West) and on the right between the Western and Central populations (T_Western-Central).

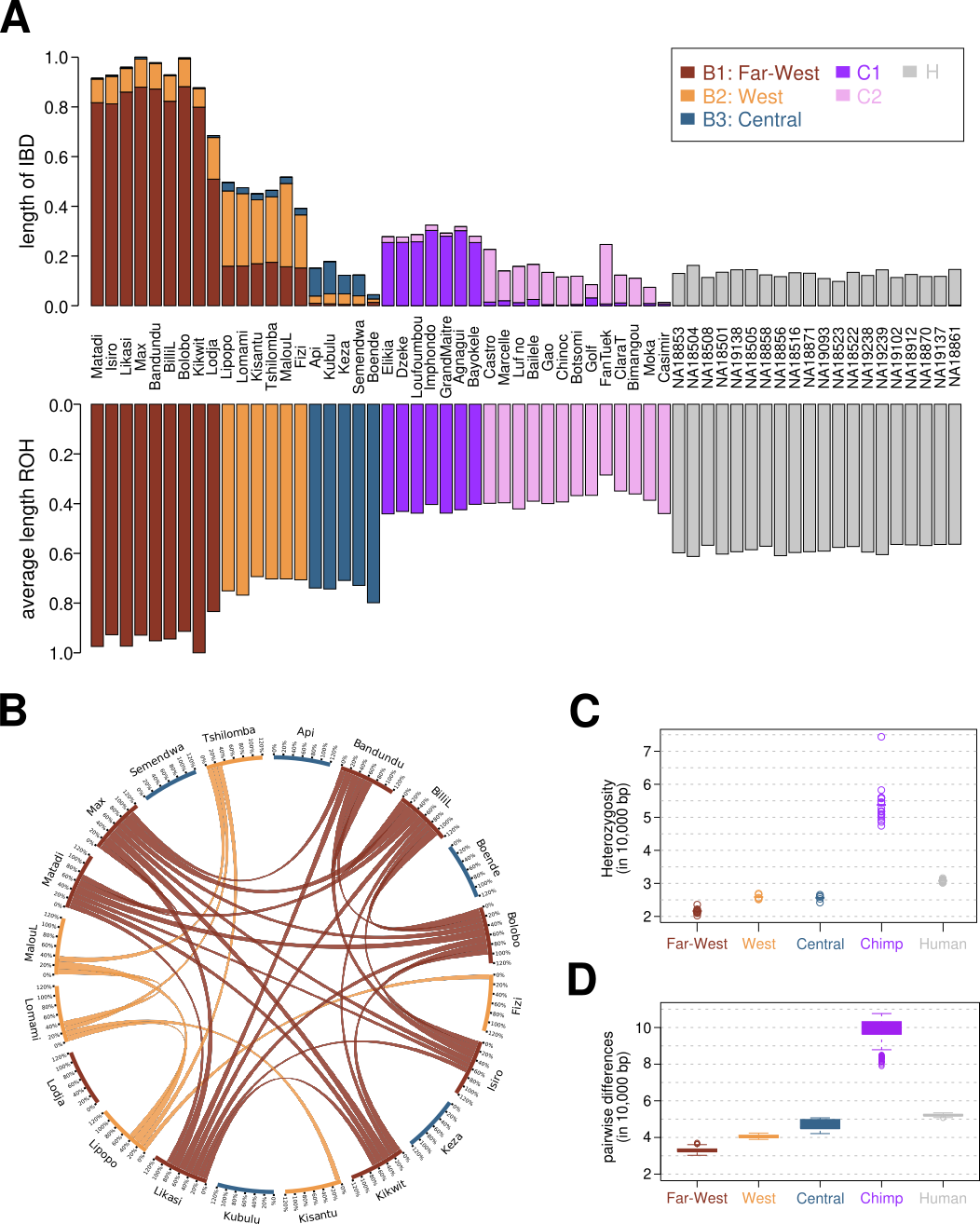

**Figure S7**

Summary statistics for each individual (A-B) and for each population/group within bonobos, and for central chimpanzees and humans Yoruba (C-D). **A.** Total length of IBD fragments per individual (upper bars) and ROH (upper bars) normalized by the maximum length observed in the sample. Each individual is represented in a vertical bar, sorted by speciess. Each segment of the vertical bar represents the length of IBD fragments shared between this individual and any individual belonging to the group of the corresponding color. The colors of bars of the ROH represent the group each individual belongs to. **B.** Inferred relatedness within the bonobo samples. Individuals are placed on the outer circle, where colors represent the group each individual belongs to. Ribbons connect pairs of individuals with a K1 estimate of at least 10%. Distribution of heterozygosity for each individual (**C**) and distribution of average pairwise differences between individuals (**D**) for the three bonobo populations, chimpanzees and humans.

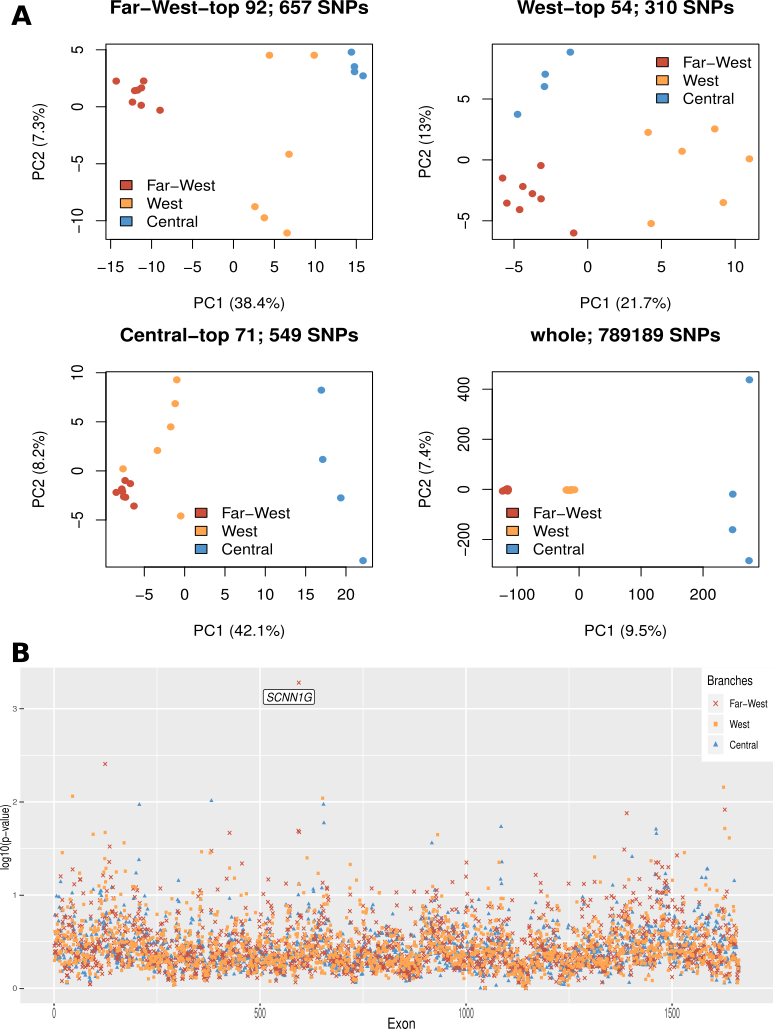

**Figure S8**

**A.** PCA of SNPs in exons with the strongest genome-wide signatures of selection (exon-wise adjusted p-value < 0.05 in GRoSS), for Far-West(B1), West(B2) and B3(Central(B). For comparison, PCA of SNPs in the whole exome (bottom right). **B.** Manhattan plot, -log10(p-values) for GRoSS per exon on chromosome 16, which contains the highest genome-wide p-value in the Far-west population (in the gene *SCNN1G*). The structure of the data, where each exon has one single p-value, results in individual low p-values rather than clear peaks.

**Table S1.** F_ST_ values among groups. **A.** Bonobo groups in our data (B1: Far-West, B2: West and B3: Central), **B.** Chimpanzee subspecies (Prado-Martinez et al. 2013) and **C**. Three continental 1KG (1000 Genomes Project Consortium, 2010) human groups, considering two populations per continent (YRI and LWK in Africa, FIN and TSI in Europe, CHB and JPT in Asia). We present one F_ST_ value within each continent, and the average of four F_ST_ pairwise population comparisons between continents.

| **A.** | |  |  | **B.** | |  |  |  | **C.** | |  |  |
| --- | --- | --- | --- | --- | --- | --- | --- | --- | --- | --- | --- | --- |
| Bonobo | Far-West | West |  | Chimpanzee | Nig-Cam | Eastern | Central |  | Human | Africa | Europe | Asia |
| West | 0.093 | - |  | Eastern | 0.163 | - | - |  | Africa | 0.013 | - | - |
| Central | 0.145 | 0.088 |  | Central | 0.166 | 0.122 | - |  | Europe | 0.136 | 0.018 | - |
|  |  |  |  | Western | 0.18 | 0.234 | 0.227 |  | Asia | 0.160 | 0.104 | 0.012 |

**Table S2 from 7. GRoSS results: SNPs with p-value < 0.05 after multiple testing correction and significant regions in each population with frequencies and annotations (extended file as an attachment).**
